# Exploring *Leishmania*-Host Interaction with Reactome, a Database of Biological Pathways and Processes

**DOI:** 10.1101/2021.03.23.436718

**Authors:** Julieth Murillo, Bijay Jassal, Maria Adelaida Gómez, Henning Hermjakob

**Affiliations:** CIDEIM - Centro Internacional de Entrenamiento e Investigaciones Médicas, Cali, Colombia; Pontificia Universidad Javeriana, Cali, Colombia; Ontario Institute for Cancer Research, Toronto, Canada; Universidad Icesi, Cali, Colombia; European Molecular Biology Laboratory, European Bioinformatics Institute (EMBL-EBI), Wellcome Genome Campus, Hinxton, Cambridgeshire, UK

**Author notes:** Address correspondence Henning Hermjakob, Maria Adelaida Gómez.

**Keywords:** *Leishmania*, Reactome, pathway, ORA, enrichment

## Abstract

Leishmaniasis is a parasitic disease with a wide range of clinical manifestations. Multiple aspects of the *Leishmania*-host interaction, such as genetic factors and modulation of microbicidal functions in host cells, influence pathogenesis, disease severity and treatment outcome. How do scientists contend with this complexity? Here, we work towards representing detailed, contextual knowledge on *Leishmania*-host interactions in the Reactome pathway database to facilitate the extraction of novel mechanistic insights from existing datasets. The Reactome database uses a hierarchy of abstractions that allows for the incorporation of detailed contextual knowledge on biological processes matched to differentially expressed genes. It also includes tools for enhanced over-representation analysis that exploits this extra information. We conducted a systematic curation of published studies documenting different aspects of the *Leishmania*-host interaction. The “*Leishmania* infection pathway” included four sub-pathways: phagocytosis, killing mechanisms, cell recruitment, and *Leishmania* parasite growth and survival. As proof-of-principle of the usefulness of the released pathway, we used it to analyze two previously released transcriptomic datasets of human and murine macrophages infected with *Leishmania*. Our results provide insights on the participation of ADORA2B signaling pathway in the modulation of IL10 and IL6 in infected macrophages. This work opens the way for other researchers to contribute to, and make use of, the Reactome database.

**Importance:** Leishmaniasis is a neglected disease infectious disease which affects more than 1.5 million people annually. Many researchers in the field apply -omic technologies to dissect the basis of clinical and therapeutic outcomes and access drug targetable features in the host-parasite interaction, among others. However, getting mechanistic insights from -omics data to such end is not an easy task. The most common approach is to use the -omics data to inquire pathways databases. The retrieved list of pathways often contains vague names that lack the biological context. In this study, we worked to create the *Leishmania* infection pathway in the Reactome database. With two practical examples from transcriptomics and microarray data, we demonstrated how this pathway facilitates the analysis of such data. In both datasets, we found a common mechanism of IL10 and IL6 production that the authors did not advert in their previous analysis, providing proof-of-principle of the tool’s enhanced potential for knowledge extraction. *Leishmania* infection pathway is in its first version, and must be expanded to cover the current knowledge base of the *Leishmania*-host interaction. We strongly encourage contributions from domain experts for the completion of *Leishmania* infection pathways.

## Introduction

The interaction between a parasite and its host is a fight for dominance. Who wins? How does disease progress? A multitude of factors determine the course of infection. These depend upon both the status of the host and the parasite. Omics data shed light on the multitude of activated mechanisms within the host-parasite interactome, by providing long lists of differentially expressed (DE) molecules. However, such data is a means for generating knowledge, not an end. The ultimate goal is to interpret this data to build a mechanistic understanding of the interactions at hand. This exercise is made tractable by the existence of pathway databases that curate and organize current knowledge (1–3). Typically, the databases are used to find known biological processes that could underlie the data. A common methodology is Over-Representation Analysis (ORA). This takes the set of DE genes from the data, and iteratively compares them to the set of genes involved in each separate pathway in the database. It uses the overlap between these two sets to predict the statistical likelihood of the biological pathway being represented in the data (4). Mechanistic hypotheses on the processes underlying the data are then proposed by the researcher, based on the ORA results.

Biological process labels within a database often lack context (e.g. ’immune system’). Does the process occur within one particular cell type, or more? Across species? In a diseased organism? In the context of a pathogen-host interaction? It is difficult to build detailed hypotheses from such labels using ORA, or indeed other analytical approaches.

The Reactome database builds a hierarchy of abstractions into which the observed features of any biological process can be incorporated. At the top level of the hierarchy, high level characteristics are represented: Is it a disease? Is it infectious or metabolic? At lower levels, features such as specific, temporally-ordered sequences of cellular processes are represented (3). Choices such as how many levels of abstraction to include, and what each should represent, depend to some extent upon the expertise of the curator. Therefore, it is critical that the curator has expert domain knowledge or that they collaborate with an appropriate expert in the field.

The purpose of this work was to add representative features and variability of the *Leishmania* spp.-host interaction into the Reactome database. In the terminology of the database, this representation is known as a ‘pathway’. Our pathway is sufficiently flexible as to allow for expansion and revision as new results are published. It incorporates detailed information on biological processes known to be activated during the *Leishmania*-host interaction. In particular, we focused on those processes correlated with the outcome of infection.

We explicitly demonstrate the utility of our database curation. We took two existing datasets to which ORA or manual revision of the literature was previously applied. With our expanded database, we uncover new mechanistic insights. In the long term, we hope that the research community will be able to use our pathways as a source of primary consultation, and as a curated database for functional and mechanistic interpretation of new data derived from -omics technologies, functional tests, or *in-silico* experiments. We believe that this pathway will enable fast and curated access to the integrative mechanisms of importance in leishmaniasis.

## Results

### Abstracting the top-level pathway: phagocytosis, killing mechanisms, cell recruitment and responses favoring *Leishmania* parasites

The curation of a pathway in Reactome starts by structuring pathways into hierarchical sections. The parent *Leishmania* infection pathway was structured into four subpathways describing the major processes involved: phagocytosis, killing mechanisms, cell recruitment and *Leishmania* parasite growth and survival. For each subpathway, there is extensive literature on the specific host responses to *Leishmania* infection, and their implication in the outcome of infection (5–7).

*Leishmania* parasites are transmitted to the host through the bite of a sand-fly that injects the motile promastigote form into the dermis of humans and other warm-blooded animals. Therein, the parasite interacts with the host cell(s) to establish the intracellular niche, where it will adopt the amastigote form (8). Paradoxically, macrophages, professional phagocytes of the innate immune system, are the main host cells for *Leishmania*. The first interaction of the parasite with the host cell is crucial to the outcome of the infection (5). The type of phagocytic receptor or pattern recognition receptor stimulated might influence the signaling cascade(s) that will trigger or inhibit cellular mechanisms involved in parasite killing or permissiveness for infection. Deregulated immune responses contribute to pathology (9). Pro and anti-inflammatory mediators must be expressed at the “right” time and in the “appropriate” magnitude, in order to have a healing response (9). This discussion motivates our abstraction of existing knowledge into the categories of phagocytosis, killing mechanisms, cell recruitment (pro-inflammatory response), and *Leishmania* parasite growth and survival (figure 1).

**Figure 1:**
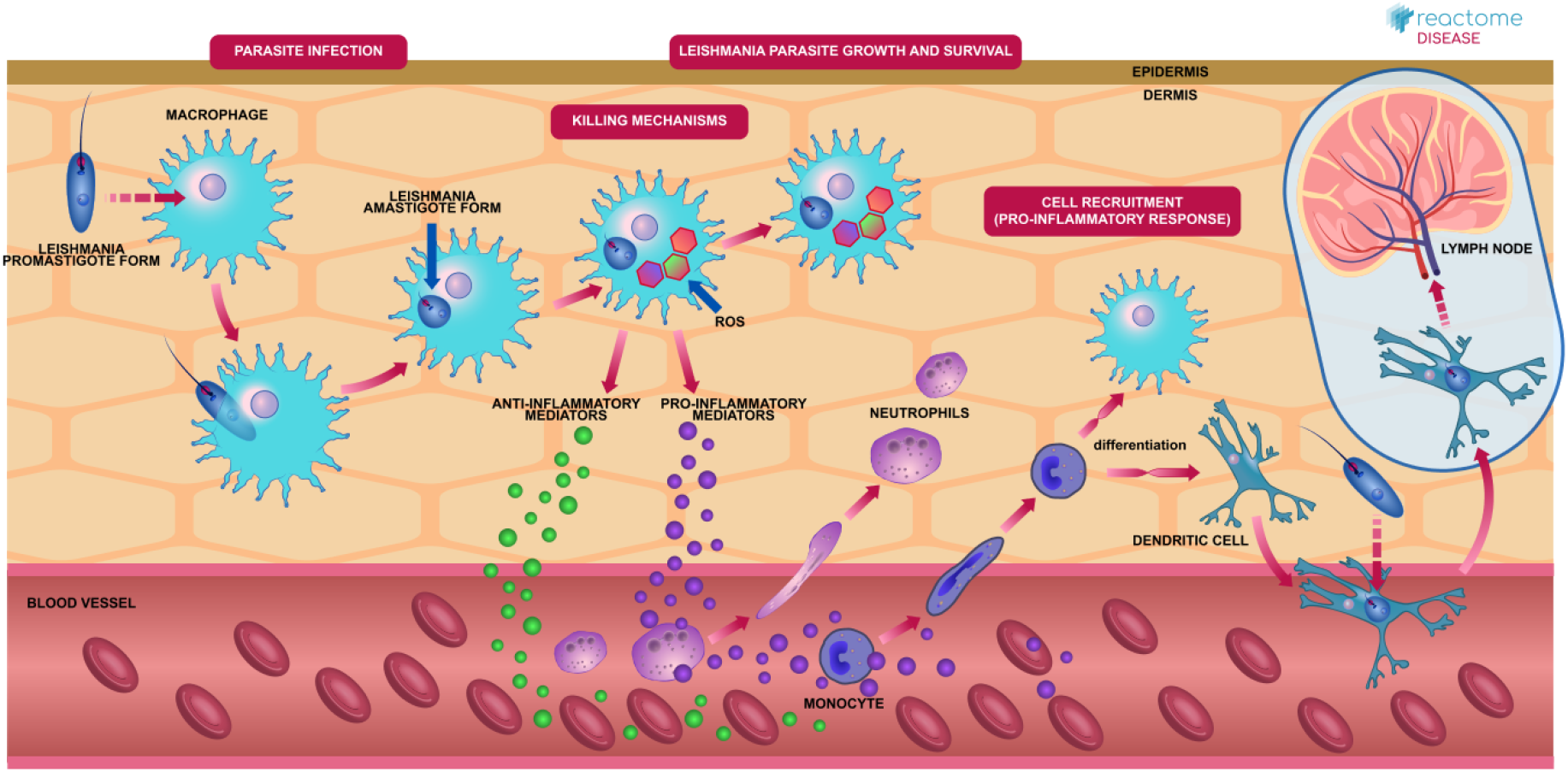
Textbook-style diagram representing the top-level pathway “*Leishmania* infection”. The major steps occurring in the dermis were compartmentalized into four categories: phagocytosis, killing mechanisms, cell recruitment, and *Leishmania* parasite growth and survival. On the webpage (https://reactome.org/PathwayBrowser/#/R-HSA-9658195), the magenta rectangular labels are interactive and take the user to the content of each subpathway.

### *Leishmania* infection pathways: From a sketch on paper to Reactome database

Reactome’s curation tool is a graphical user interface (GUI), that connects to its central database with which new information can be added to existing or new pathways. We started with a sketch of the overall pathways we wanted to curate for the first version of the “*Leishmania* infection pathway”. Then, we identified what molecules/entities and reactions already existed in the database and which ones needed to be added. Similarly, we accounted for the molecular interactions that were already described in existing Reactome pathways. If new reactions were required, they were created from scratch, supported by literature references. Table 1 summarizes these metrics.

**Table 1:**
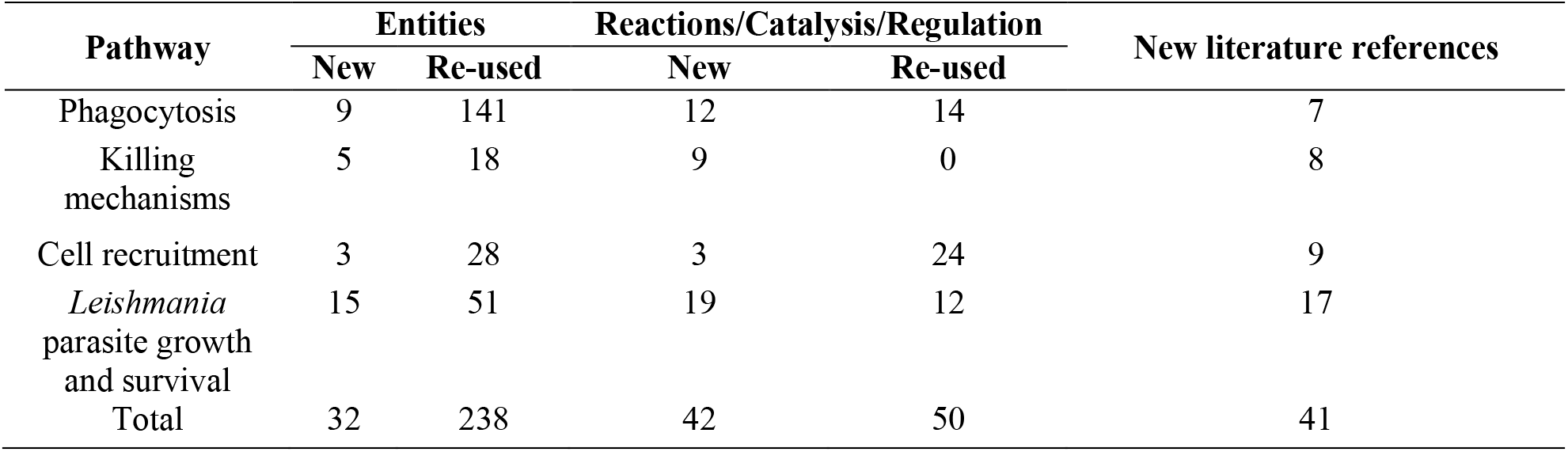
Curation and annotation metrics for the creation of *Leishmania* infection pathways in Reactome.

| Pathway | Entities |  | Reactions/Catalysis/Regulation |  | New literature references |
| --- | --- | --- | --- | --- | --- |
|  | New | Re-used | New | Re-used |  |
| Phagocytosis | 9 | 141 | 12 | 14 | 7 |
| Killing mechanisms | 5 | 18 | 9 | 0 | 8 |
| Cell recruitment | 3 | 28 | 3 | 24 | 9 |
| *Leishmania* parasite growth and survival | 15 | 51 | 19 | 12 | 17 |
| Total | 32 | 238 | 42 | 50 | 41 |

### Structuring the lowest-level pathways : signaling cascades traceable from a membrane protein to the production of effector molecules

As part of the host response against *Leishmania*, many signaling cascades are modulated (activated or inactivated). Once a cellular/membrane receptor is stimulated, the downstream signal transduction can result in the activation of many molecules with different effects in the system (e.g. interleukins inducing the polarization of several types of T-cells, or chemokines mediating the recruitment of immune cells to different tissues, among others). That is why general pathway labelling such as “TNF-signaling”, might not be informative enough, and can allow for erroneous biological interpretations if the gene lists contained within enriched pathways are not carefully analysed. There could be many regulatory processes that favor one direction rather than another in a specific signaling pathway (as it can be noticed in TNF-pathway in Reactome, R-HSA-75893 https://reactome.org/PathwayBrowser/#/R-HSA-75893 and KEGG, hsa04668-https://www.genome.jp/kegg-bin/show_pathway?hsa04668). Therefore, we structured the lowest-level pathways in each of four subpathways, starting off from an activated membrane protein (e.g., receptors, ion channels or enzymes). This was followed by inclusion of signalling and accessory molecules and finished with synthesis of effector molecules that are consistent with the overall biological processes underlying the subpathway (e.g., reactive oxygen species –ROS-for “killing mechanisms”).

For this first version of the *Leishmania* infection pathway, we chose the membrane proteins FCGR3A, FZD7, P2RX4, P2RX7, ADORA2B, and CD163 and their downstream signaling cascades. Although additional membrane molecules are known to participate in the initial macrophage-*Leishmania* interaction, such as complement receptors, toll-like receptors -TLRs-, or chemokine receptors among others (10–12), we selected less “classical” membrane molecules to increase the breath of mechanistic interpretation of host-*Leishmania* –omic datasets. Shown in figure 2 are the structures of the four subpathways, each of which contains a set of reactions. Supporting references evidencing their relevance in the context of leishmaniasis will be discussed below.

**Figure 2:**
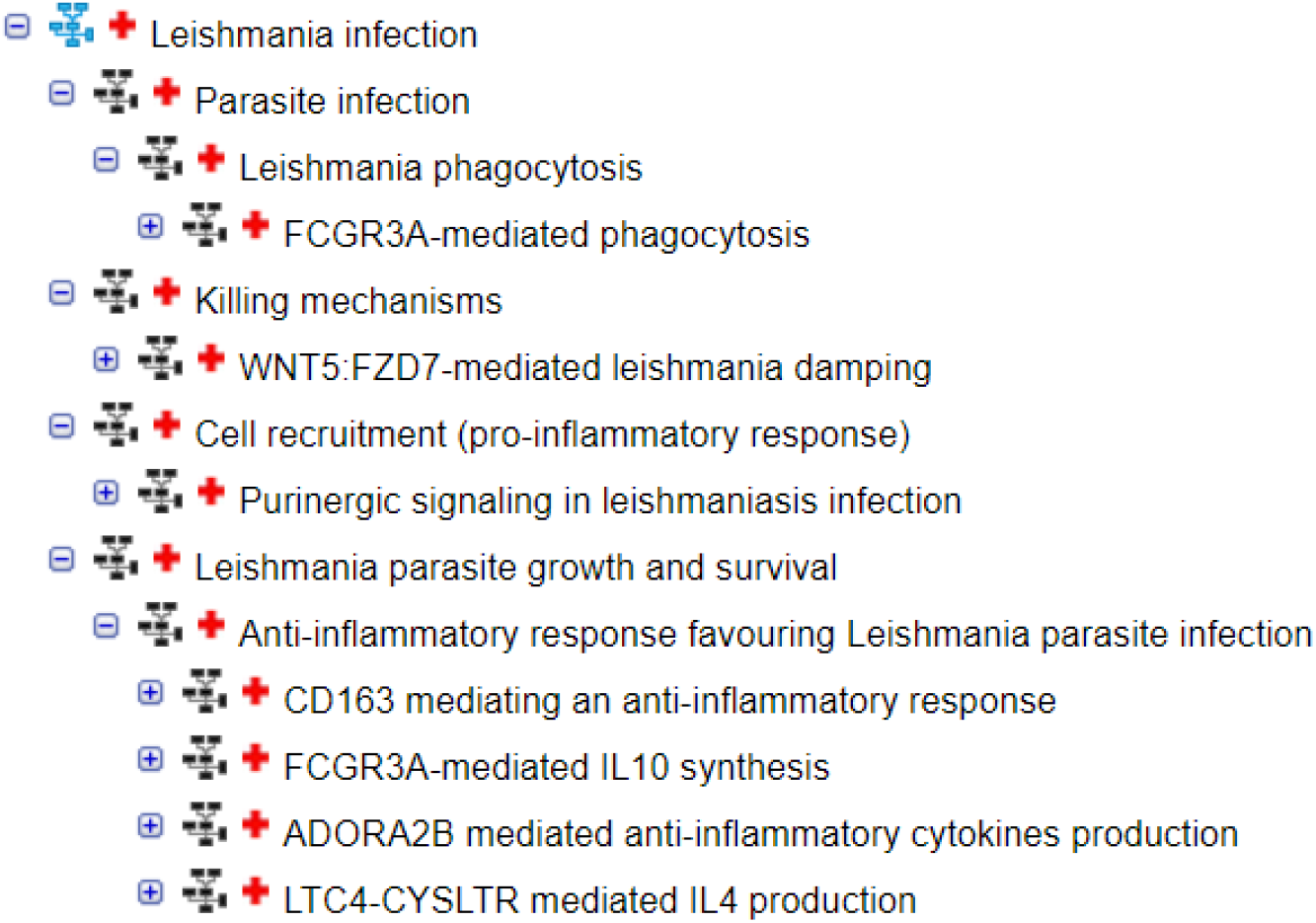
Hierarchical structure of the *Leishmania* infection pathway showing the four sub-pathways and their contents. Each indentation introduces a new subpathway; as an example, “parasite infection” is the parent pathway for the subpathway “*Leishmania* phagocytosis”, which, to date only contains *FCGR3A-*mediated phagocytosis.

I. **Parasite Infection/ *Leishmania* phagocytosis**: The phagocytosis subpathway was built to account for the different types of phagocytic receptors that *Leishmania* parasites can utilize for their entry into host cells. We started by adding FCGR3A-mediated phagocytosis. To build it, we referenced the pre-existing Reactome pathway “FcG receptor (FCGR) dependent phagocytosis” (https://reactome.org/content/detail/R-HSA-2029480). Overall, 14 reactions were re-used from this pathway while 12 new reactions were created to represent phagocytosis in the context of *Leishmania* infection (table 1). The starting point was the binding of immunoglobulin G antibodies (IgG) to either an unknown *Leishmania* amastigote (abbreviate as: Lma) surface molecule or the glycoinositol phospholipid -GIPL (13, 14), as shown in figure 3a. These interactions correspond to binding reaction types in Reactome, with the product of binding being a complex comprising the inputs. The complexes “IgG:Lma surface” and “IgG:GIPL” represent the opsonization of the *Leishmania* amastigote by the antibody IgG, during a “second round” of host-parasite contact where the proliferative form and infective form in the host is the amastigote. We assumed the same course when opsonization occurs via these known, or other unknown molecules. Therefore, we collated the two complexes into one entity, which in Reactome is represented by the defined set “IgG:Lma antigens”. From here, downstream reactions continue towards the activation of actin filaments that then continue to the formation of the phagocytic cup. The overall diagram depicting each step of the pathway can be accessed through this link https://reactome.org/PathwayBrowser/#/R-HSA-9664422&PATH=R-HSA-1643685,R-HSA-5663205,R-HSA-9658195,R-HSA-9664407,R-HSA-9664417. A close-up depicting a portion of the pathway is found in figure 3A.

**Figure 3:**
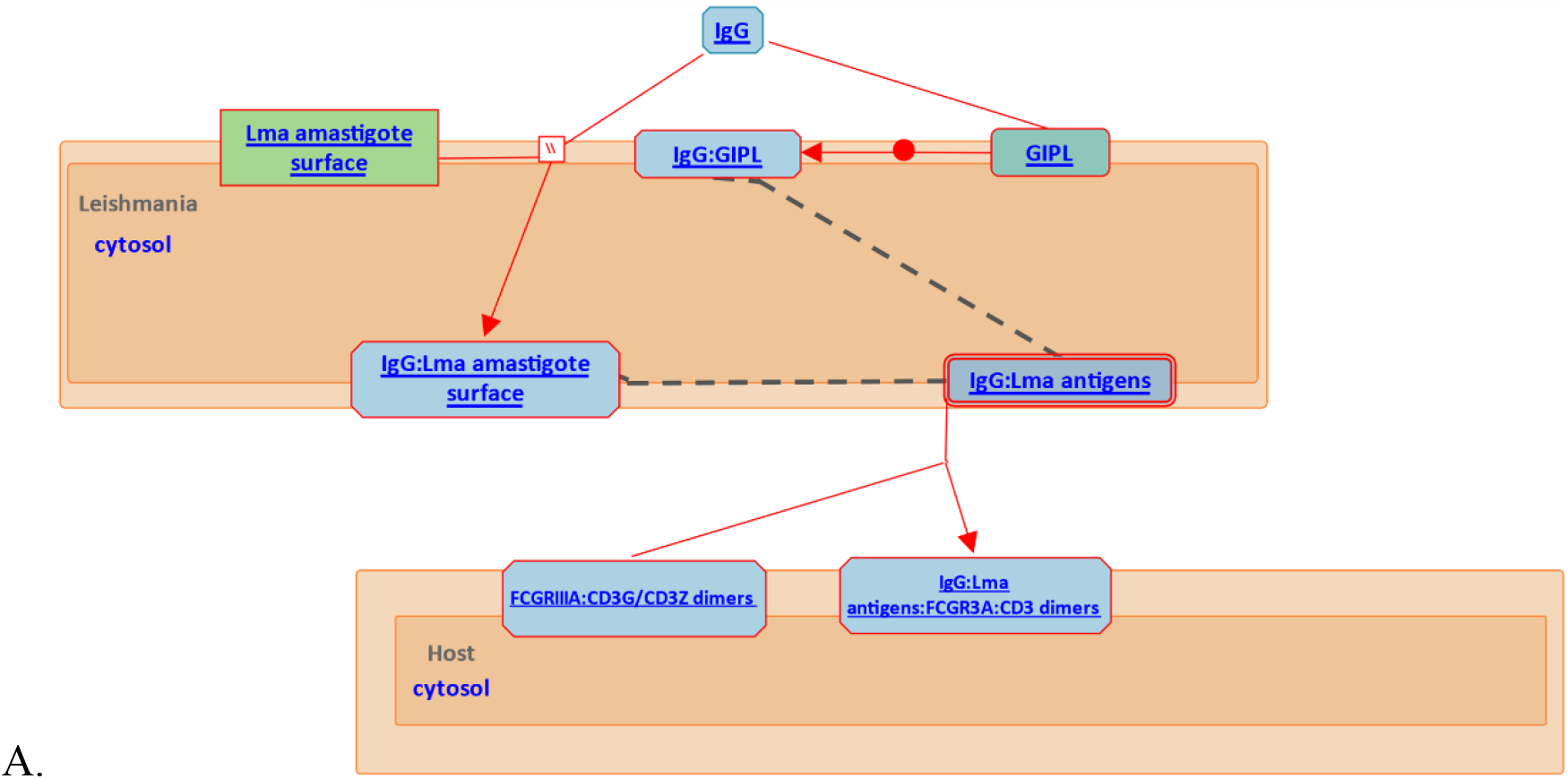
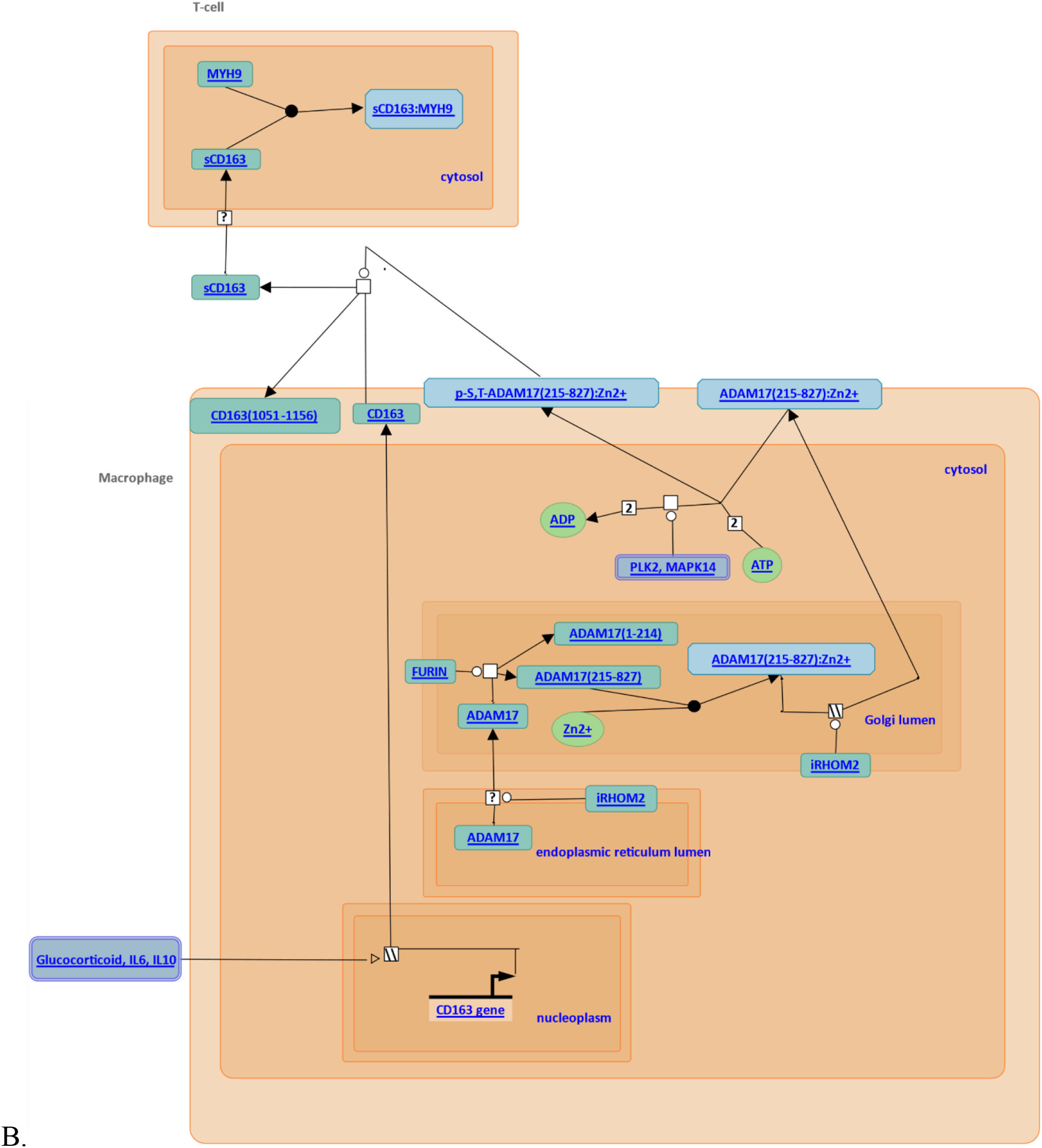
Standard graphical representation of pathways in Reactome. **A**. Fragment of the diagram for FCGR3A-mediated phagocytosis. Shown are the reactions corresponding to the parasite opsonization process by IgG. Parasitic components are highlighted in red. **B**. CD163 mediating an anti-inflammatory response. Each diagram shows the participating entities in the granularity of chemical compounds (green ovals), proteins (green rectangles), complexes (blue rectangles) and sets (blue rectangles with a double border). The arrangement of the entities in the reactions can be easily followed on the web page, by clicking on the arrows that connect adjacent steps. The route of entry into the macrophage can affect the fate of *Leishmania* parasites (5). We expect to incorporate the internalization processes that are mediated by other receptors into the *Leishmania* phagocytosis subpathway. This includes complement receptors (CR3 and CR1), mannose receptor-MR, and fibronectin receptors-FNRs (5). Similarly, the “Parasite infection” subpathway will be populated with the steps that describe the maturation of the phagocytic cup, and so on.
II. **Killing mechanisms**: this subpathway was designed to contain the signaling cascades that converge in the production of antimicrobial molecules in the context of leishmaniasis. We started by curating the activation of the receptor Frizzled-7 (FZD7) by the ligand WNT5 and its downstream cascade. To build this pathway we reviewed publications that contain the original experimental data used to determine the reactions details (15–19). WNT5 is known for being a highly specific regulated gene in response to microbial infection (20–22) including leishmaniasis (23), where it seems to be involved in mechanisms that dampen the parasite load within the macrophage. Complementary, FZD7 acts as a receptor of WNT5 which, upon binding, is implicated in the initiation of the non-canonical WNT pathway that leads to re-organization of the cytoskeleton to allow a process called planar cell polarity (PCP) (22). The activation of the WNT5:FZD7 non-canonical signaling cascade that drives PCP is being studied for its involvement in inflammatory responses (24). Treatment of RAW264.7 macrophages with recombinant WNT5 induced NADPH oxidase-mediated ROS production, which has been suggested to contribute to the macrophage control of *L. donovani*. Consequently, detailed understanding of how the WNT signaling network defines host responses to infection could be important to identify new potential therapeutic targets (22). We represented in 9 reactions, the activation of FZD7 by the WNT5 ligand, resulting in the production of ROS (table 1). Unlike the phagocytosis pathway, these reactions correspond to a host’s response to the infection, even if no parasite components are depicted in the diagram (found at https://reactome.org/PathwayBrowser/#/R-HSA-9673324&PATH=R-HSA-1643685,R-HSA-5663205,R-HSA-9658195,R-HSA-9664420). In future versions we will incorporate the cross talk with signaling cascades, like TLR-signaling, that activate antimicrobial functions and synthesis of antimicrobial molecules.
III. **Cell recruitment:** this subpathway was aimed at bringing together signaling pathways that converge in the induction of gene expression and synthesis of chemokines and pro-inflammatory cytokines. It is known that a proinflammatory response early in the infection enhances host cell microbicidal mechanisms (25). However, the recruitment of inflammatory cells to the site of infection, once the parasite load has been controlled, transforms the course of infection and can lead to immunopathology (9). Therefore, it is important to curate and represent specific pathways that have shown to be activated upon *Leishmania* infection, resulting in the production of pro-inflammatory mediators. The first specific mechanism we curated was the activation of the purinergic receptors P2RX4 and P2RX7. The liberation of ATP normally occurs in tissues facing stressful stimuli such as infection (26). Binding of ATP to purinergic receptor activates the inflammasome leading to subsequent activation of interleukin 1 beta-IL1β, which promotes the recruitment and activation of macrophages (27). We represented this process in 27 reactions (table 1), that included a regulatory step mediated by NTPDase1 and NTPDase5, which reduces ATP to adenosine (28). The molecular diagram can be found at https://reactome.org/PathwayBrowser/#/R-HSA-9664424&SEL=R-HSA-9660826&PATH=R-HSA-1643685,R-HSA-5663205,R-HSA-9658195. There are many other pathways promoting cell recruitment as a response to *Leishmania* infection with different consequences for the parasite and the host (29). In future expansions of this subpathway, it would be possible to highlight cross talk between different cascades that target the same effector molecules.
IV. ***Leishmania* parasite growth and survival:** this subpathway covers the host responses that favor intracellular parasite survival, and the mechanisms used by the parasite to hijack host cell functions. To survive as an intracellular parasite, *Leishmania* evades the activation of host cell microbicidal machineries. Many mechanisms facilitate this purpose. On the host side, the production of anti-inflammatory mediators often occurs alongside the repression of expression of antimicrobial molecules, together with the recruitment of regulatory immune cells (e.g., regulatory T-cells). From the parasite side, inactivation of host molecules through mechanisms such as cleavage or activation of phosphatases are part of its repertoire (9). Induction of anti-inflammatory molecules was the first mechanism that we curated, compiling the steps that describe the cleavage of the membrane protein CD163, the activation of the receptors FCGR3A and ADORA2B, and ending with the corresponding production of the known anti-inflammatory molecules sCD163, IL4, and IL10, as well as the dual functioning IL6. Macrophages infected with *L. amazonensis* or *L. donovani* strongly express the membrane protein CD163 (30–32), and soluble CD163 (sCD163) has been proposed as biomarker of visceral leishmaniasis. The hypothesis of the association between sCD63 and an anti-inflammation status is that it interferes with the proliferation of T-cells (33, 34). sCD163 is formed from the increased shedding of CD163 mediated by the metalloprotease ADAM17 (35, 36). Posteriorly, it might translocate to the cytoplasm of T-cells (through an unknown mechanism) where it binds with a protein involved in the proliferation process (33, 34). In “CD163 mediated anti-inflammatory responses” we represented, in 9 reactions, the production of sCD163 including the steps that precede the activation of ADAM17. Additionally, we included the positive regulation of glucocorticoids, IL6 and IL10 on CD163 gene expression (37–40). In figure 3B we show the molecular diagram depicting the pathway. IL-10 is an important immunoregulatory cytokine produced by many cell populations; in macrophages it is induced after the stimulation of TLRs, FCG receptors or by TLR-FCGR crosstalk (41). Classically, its function is considered to be the limitation and termination of inflammatory responses and the regulation of differentiation of several immune cells (42). In the context of leishmaniasis, IgG-opsonized amastigotes have been shown to induce IL10 production through FCGRs, which in turn suppresses the killing mechanisms in phagocytic cells (14). We represented, in 21 steps, the activation of FCGR3A that leads to the activation of the transcription factor CREB1, ending with the production of IL10. Finally, we curated ADORA2B-mediated anti-inflammatory responses. ADORA2B is a receptor for the ribonucleoside adenosine. Its activation leads to the production of anti-inflammatory cytokines which have been shown to favor *Leishmania* infection and survival (43–45). Apparently, this pathway exerts an opposing/regulatory response to the purinergic signaling pathway. The blockade in the production of pro-inflammatory cytokines may come with the inhibition of killing mechanisms (46). We represented this pathway with 6 reactions, starting with the binding of adenosine to ADORA2B, and ending with synthesis of IL6. Both FCGR3A and ADORA2B signaling pathways activate transcription factors, generating a positive feedback loop for transcription of more anti-inflammatory cytokines (47, 48). In future versions, these reactions might be incorporated, as well as other pathways leading to the synthesis of other anti-inflammatory mediators known to be induced during the *Leishmania*-host interactions. Moreover, other mechanisms that favor the persistence of *Leishmania* parasites must be added into new subpathways (e.g., Polyamine synthesis). The representation of these pathways in Reactome is available in the following link: https://reactome.org/PathwayBrowser/#/R-HSA-9662851&PATH=R-HSA-1643685,R-HSA-5663205,R-HSA-9658195,R-HSA-9664433.

### *Leishmania* infection pathways enhancing transcriptome data analysis

The first version of *Leishmania* infection pathways was released in March 2020. All released data can make use of Reactome pathway analysis tools. We used the ORA tool to test the impact of context-dependent labels on published datasets (prior march-2020) that explored the *Leishmania*-host interaction.

In 2015, Dillon *et al*. (49) explored the early response of macrophages to infection with *L. major*, using RNA-seq. With the differentially expressed (DE) gene list (>|2|-fold, uninfected macrophages versus infected macrophages at 4 hours post-infection), they performed ORA against the curated pathways in the KEGG database. For the up-regulated genes, the results included cytokine-cytokine receptor interactions, TNF-signaling pathway and NFkappa B-signaling pathway, among other pathways. An overall interpretation of these enriched pathways may suggest that the early infection of macrophages by *L. major* leads to induction of pro-inflammatory responses; as an example, TNF-signalling as represented in KEGG (PATHWAY: hsa04668), classically leads to the recruitment of inflammatory cells. However, transduction of the signal may induce the activation of factors that contribute to opposite responses, such as tissue regeneration (with VEGF and EDN1) (50, 51). Interestingly, among their gene-specific analyses, Dillon *et al*. identified a set of genes involved in anti-inflammatory responses (*Csf1, Csf3, Il10, Il11r, Il1rn, Socs3, Hmox1, Egfr* and, *Vegf*). However, the mechanisms that lead to production of these effector molecules, or how they contribute to achieve or maintain the underlying immune status couldn’t be inferred from the KEGG pathway analysis. To overcome this gap, we implemented ORA from the same DE gene list (supplementary table S1) in Reactome. Among enriched pathways (table S1), were some of the *Leishmania* infection subpathways. As reported by Dillon and colleagues, FCGR3A subpathway (figure 4A) was found downregulated. Interestingly, the most over-represented pathway was the ADORA2B mediated anti-inflammatory cytokine production (Fold change-FC = 3.25). This pathway contributes to the expression of IL6 (FC = 7.68) and IL10 (FC = 17.05), through the activation of the transcription factor CREB (FC = 2.75) (figure 4B). This pathway, simultaneously, leads to the activation of killing mechanisms, as well as to the production of IL10. Neither authors nor we found pathways involved in the former. Therefore, this suggests an alternative mechanism mediated by ADORA2B, that leads to production of IL10, and does not induce parallel pro-inflammatory consequences or the activation of antimicrobial molecules.

**Figure 4.**
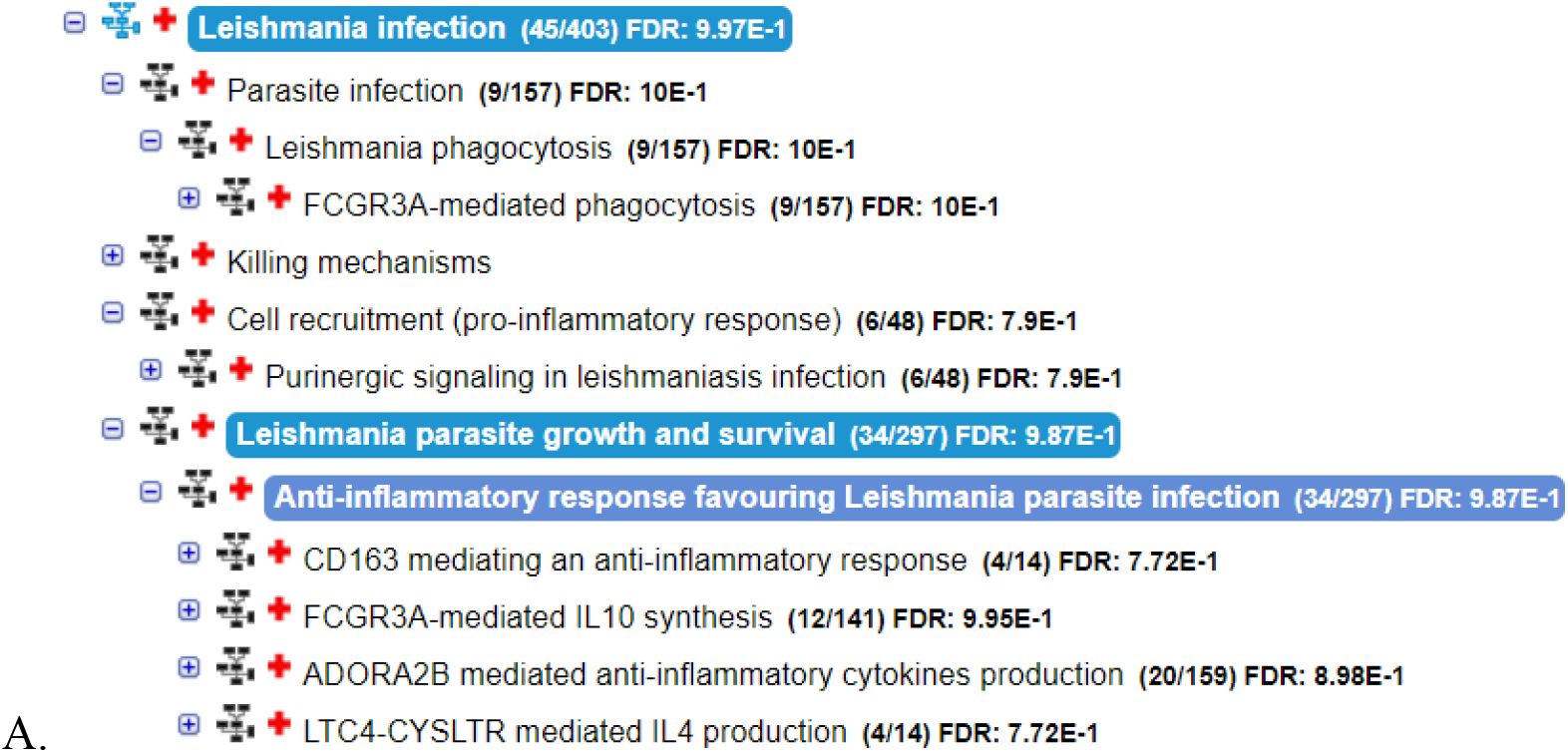
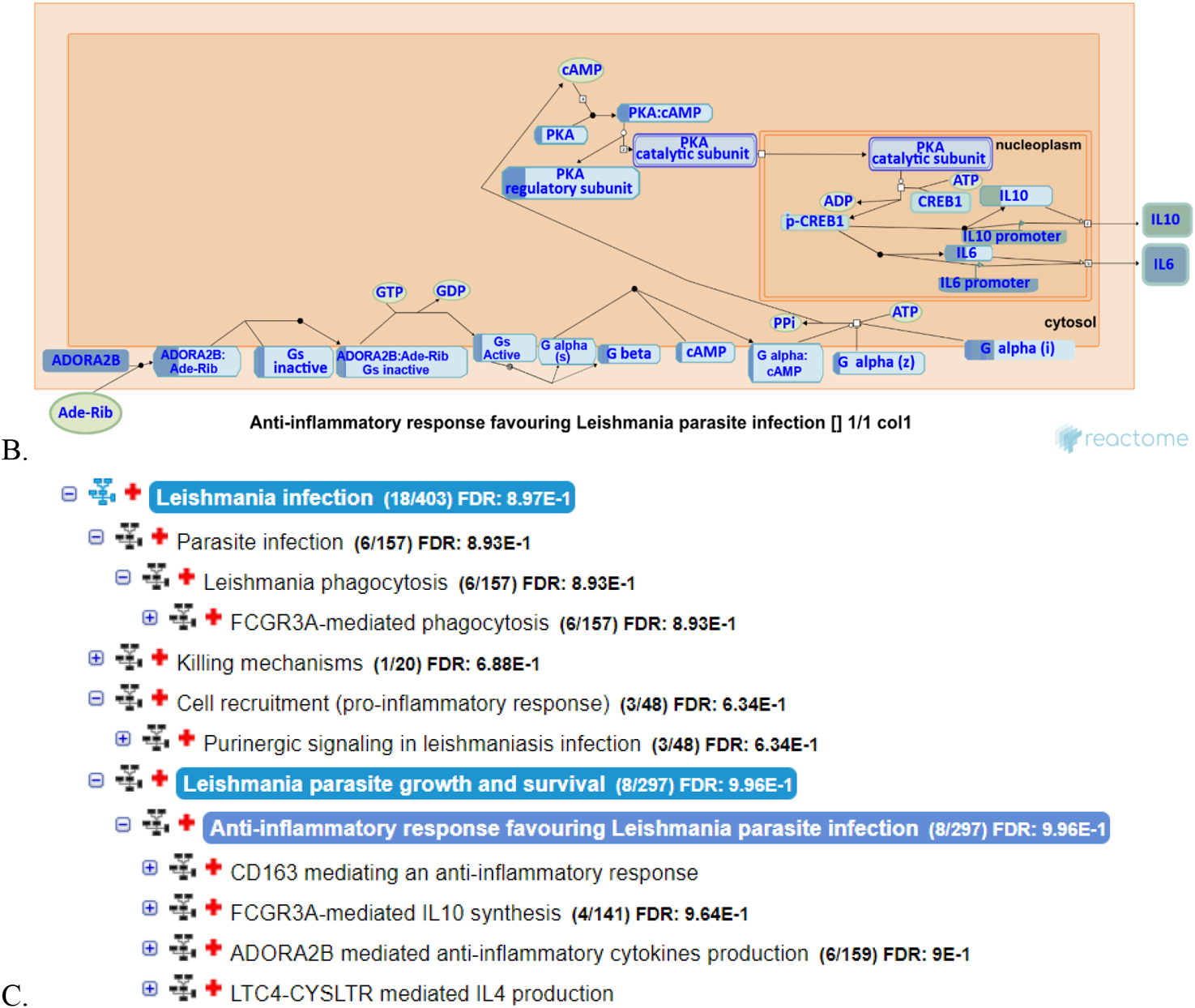
Results of applying ORA in Reactome, on the datasets generated by Dillon *et al*. and Gregory *et al*. A. Number of genes found in each subpathway from Dillon’s dataset, with the associated false discovery rate (FDR) B. ADORA2B pathway enriched in Dillon’s dataset shows upregulated genes involved in ADORA2B signaling cascade leading to the production of IL6 and IL10. C. Number of genes found in each subpathway from Gregory’s dataset, and associated FDR.

Complementary, we analyzed the microarray data from Gregory *et al*. (52) which represents the transcriptomic response of murine macrophages to infection with *L. major* and *L. donovani*. The authors stated that there were few differences in the number of genes and the magnitude of the expression with both species. Like Dillon’s dataset, the ADORA2B subpathway was enriched among upregulated genes (figure 4C and supplementary table S2). These results highlight the identification of ADORA2B-pathway driving anti-inflammatory cytokine production, consistently activated in macrophages infected with different *Leishmania* species. These results substantiate the usefulness of the *Leishmania* infection pathways for gaining mechanistic insights from new and previously published data. In expanded versions of this pathway, secondary analyses like the one we performed, would shed light on response patterns against the infection with different *Leishmania* species, as well as species-dependent responses.

## Discussion

Over-representation analysis (ORA) has become one of the standard methods for extracting mechanistic information from -omics datasets. As part of the workflow, ORA matches the omics data with molecular data curated in pathway databases. As a result, it gives a list of pathway names in which the genes are known to be involved. Mechanistic insight must then be built from a list of labels, a problem made much harder when labels lack biological context. There are several specific cases in the literature, relating to *Leishmania*-host interaction, where this has proven to be an issue (49, 53, 54). In this work, we addressed this by creating leishmaniasis-context pathways in the Reactome database. We created four subpathways, labelled as: *Leishmania* phagocytosis, killing mechanisms, cell recruitment (pro-inflammatory response), and *Leishmania* parasite growth and survival. Inside each, we labeled the pathways according to the receptor or membrane protein directing the signaling cascade. This structure facilitates the generation of high-level mechanistic insights from low-level processes highlighted by ORA, which are sensitive to particular experimental contexts (e.g. the source of the biological sample the data was derived from).

Manual literature search is often required for in depth interpretation of the outcome of ORA for extracting biological/functional insights. However, this strategy is time consuming, prone to omissions, and certainly incompatible with unbiased exploration of novel mechanisms/functions in the data, which is the goal of –omic approaches. This is because the search is necessarily constrained to cover a small number of topics within the researcher’s area of expertise, and it is unachievable to manually trawl the entirety of biological literature relating to a particular gene, microorganism or disease.

We have re-analyzed two previously published transcriptomic datasets (from *Leishmania*-infected macrophages), to provide proof-of-concept of the usefulness of context-dependent databases such as the one described in this study. Results from this secondary analysis revealed the putative participation of a signaling pathway (ADORA2B mediated anti-inflammatory cytokines production) in the parasite-mediated induction of anti-inflammatory molecules.

Although *Leishmania* infection pathways in its first version is far from representing the full current knowledge about the interaction between the parasite and the host, we have shown that our database curation has led to new mechanistic insights from existing datasets. Moreover, these findings are generated from the use of a workflow that skips particular time-consuming, problematic manual curation steps, aligning the available data interpretation tools to the nature of unbiased hypothesis generation from –omics datasets.

## Materials and Methods

### *Leishmania* Infection Curation Process

The abstraction of *Leishmania* infection into a structure of pathways and reactions fulfilling the Reactome paradigm was accomplished by the Reactome working group. This consisted of a consortium of biocurators, software developers and leishmaniasis researchers. The latter selected the mechanisms of interest. The selection was based on biological pathways known to be associated with the infectious outcome, directly or indirectly. From here, domain experts and Reactome curators worked side by side to translate the selected pathways into the Reactome data structure using the curator tool version 3.3.

Reactome represents, categorizes, and annotates all known entities/molecules in each reaction. Different components can interact with each other only in ways prescribed by the Reactome data-model. For instance, the representation of an individual protein in the database requires several steps. In an example, for the protein ADAM17, the UniProt ID (P78536) is retrieved, as well as the GO cellular compartment it functions, and which species this protein belongs to (e.g., *Homo* sapiens). Note that if the represented process implies the transition of the same protein from one compartment to another, different instances of this protein must be created (e.g ADAM17 [endoplasmic reticulum], ADAM17 [golgi apparatus] and ADAM17 [plasma membrane]). However, all these instances still point to the same UniProt ID. On the other hand, if several proteins individually fulfil the same role in a reaction (e.g., phosphorylation), these are grouped together into a single entity called “*defined set*” (PKL2 [cytosol] and MAPK14 [cytosol] in the *defined set* PKL2, MAP14 [cytosol] that phosphorylates ADAM17 [plasma membrane]). The *defined set* entity is also specific to each cellular compartment. Therefore, each individual molecule within the *defined set* must have a compartment-specific representation. Otherwise, in the quality assurance procedure, this will come out as an error. Molecules of the same or different types can form complexes, for example ADAM17:Zn2+ which represents the functional form of ADAM17. Once these entities are created, the reactions in which they take part, are created. Binding is a common type of reaction in Reactome. Here, the output is a complex type entity (e.g., the interaction between sCD163 and MYH9 is represented as a binding that ends in the formation of the sCD163:MYH9 complex). If we are not at the final point of the pathway or dealing with a secondary product, the complex can participate in subsequent reactions. In that case, the reaction from which this complex came from is indicated as the “preceding event” of the subsequent reaction.

Overall, we curated *Leishmania* infection pathways following the general structure of a signaling pathway. Namely, a ligand binding to a receptor, then the stimulated receptor effecting a downstream signaling cascade, up to the activation of effector molecules (eg. cytokines, nitric oxide, etc.). However, other pathways were conceived with different starting points. For instance, the first step of the CD163 example pathway consisted of ADAM17 activation. This begins in the endoplasmic reticulum, and its maturation process follows through the Golgi apparatus until its translocation to the plasma membrane, where its phosphorylation by kinases (either PLK2 or MAPK14), activates the cleavage reaction of CD163. It is the soluble portion of CD163, sCD163, that has been found as a regulator of the inflammatory responses in *Leishmania* infection, through the inhibition of the proliferation of T lymphocytes.

Reactome’s criteria for the acceptance of a particular molecular interaction and its supporting reference have been previously explained (3, 55). Once the reactions were successfully integrated into Reactome’s central database, we reached out to an experienced researcher in leishmaniasis, Dr. David Gregory (ORCID: 0000-0001-6534-7150) (52, 56, 57) to review the curated material on the basis of his expertise in the field (52, 57–59). Only material reviewed by an independent domain expert is allowed to be published in a Reactome release. The contributions of authors and reviewers can be directly accessed through Reactome’s search interface, for example https://reactome.org/content/query?q=David+Gregory shows a detailed description of credit attribution in Reactome (60). We are highly interested in further contributions from domain experts for the completion of *Leishmania* infection pathways.

Once an orderly list of reactions was created, these were transformed according to the Reactome paradigm by using the curator tool. Some reactions were created from scratch to populate a low-level pathway, while others that were present within other pathways were duplicated (or reused) in the context of *Leishmania* infection (table 1). Within a reaction, each component was re-used if it already existed as part of a different process in the database, otherwise it was created. Reuse of existing components and reactions ensures no redundancy in the database.

Reaction details, such as input/catalyst/output molecules, reaction type, preceding reactions and experimental species, were captured during the curation process. Reactions were linked together based on preceding-following relationships to generate pathways.

## Suplementary material description

S1: It is an excel file with two tabs. The first one contains the gene input from the paper by Dillon *et al*. (49). The second one contains the output of applying ORA to that gene list.

S2: It is an excel file with two tabs. The first one contains the gene input from the paper by Gregory *et al*. (52). The second one contains the output of applying ORA to that gene list.

